# Evolutionary Rescue Over A Fitness Landscape

**DOI:** 10.1101/135780

**Authors:** Yoann Anciaux, Luis-Miguel Chevin, Ophélie Ronce, Guillaume Martin

## Abstract

Evolutionary rescue describes a situation where adaptive evolution prevents the extinction of a population facing a stressing environment. Models of evolutionary rescue could in principle be used to predict the level of stress beyond which extinction becomes likely for species of conservation concern, or conversely the treatment levels most likely to limit the emergence of resistant pests or pathogens. Stress levels are known to affect both the rate of population decline (demographic effect) and the speed of adaptation (evolutionary effect), but the latter aspect has received less attention. Here, we address this issue using Fisher’s Geometric Model of adaptation. In this model, the fitness effects of mutations depend both on the genotype and the environment in which they arise. In particular, the model introduces a dependence between the level of stress, the proportion of rescue mutants, and their costs before the onset of stress. We obtain analytic results under a strong-selection-weak-mutation regime, which we compare to simulations. We show that the effect of the environment on evolutionary rescue can be summarized into a single composite parameter quantifying the effective stress level, which is amenable to empirical measurement. We describe a narrow characteristic stress window over which the rescue probability drops from very likely to very unlikely as the level of stress increases. This drop is sharper than in previous models, as a result of the decreasing proportion of stress-resistant mutations as stress increases. We discuss how to test these predictions with rescue experiments across gradients of stress.

## INTRODUCTION

Understanding the persistence or decline to extinction of populations facing environmental stress is a crucial challenge both for the conservation of biodiversity and the eradication of pests or pathogens (Gonzalez *et al.* 2013; Carlson *et al.* 2014; Alexander *et al.* 2014; Bell 2017). In evolutionary biology, environmental stress describes any conditions in the environment that induces a reduction in individual fitness (Koehn and Bayne 1989; Bijlsma and Loeschcke 2005). Here, we will focus on the case where environmental stress causes a reduction of population mean fitness that is harsh enough to trigger a decline in abundance (Hoffmann and Parsons 1997). In such a stressful environment, if heritable variation in fitness is available or arises by mutation, adaptive evolution may allow the population to escape extinction. This phenomenon has been called evolutionary rescue (ER) (Gomulkiewicz and Holt 1995). Evolutionary rescue is of particular importance for understanding the emergence of genetic resistance to drugs or treatments in medicine and agronomy (Davies and Davies 2010).

Empirical evidence supports the idea that stress levels critically determine ER probabilities (Samani and Bell 2010; Moser and Bell 2011; Lindsey *et al.* 2013). For example, the probability that bacteria evolve antibiotic resistance (that is, the probability of avoiding antibiotic-induced extinction through ER) typically declines sharply, in a strongly non-linear way, with increasing drug concentration (Drlica 2003). Evolutionary rescue thus shifts from being highly likely to highly unlikely over a narrow window of stress levels. This critical range of stress depends on the strain, especially on its evolutionary history with respect to exposure to the stress (Gonzalez and Bell 2013). Stress level, as controlled by drug concentration, has also been shown to affect the genetic basis of resistance (e.g. Harmand *et al.* 2017), with a wider diversity of genes and alleles conferring resistance at low than at high doses. However, the underlying causes for the relationship between stress level and ER are still poorly understood. Our aim here is to derive new analytical predictions for this relationship. In particular, we want to predict the critical window of stress levels above which ER is very unlikely, allowing direct comparison with experimental data.

In the theoretical literature (reviewed in Alexander *et al.* 2014), most ER models predict that ER probability decreases with increasing stress level, measured by the decay rate of the stressed population. Indeed, a faster decay of the population leaves less time for adaptation to occur before extinction (e.g. Gomulkiewicz and Holt 1995). But beyond this direct demographic effect, stress level may also have indirect effects on ER. Indeed, a stressful environment may not only affect the demographic properties of the population, but also its rate of adaptation, by modifying the determinants of genetic variance in fitness (Hoffmann and Parsons 1997; De Visser and Rozen 2005; Agrawal and Whitlock 2010). First, the rate of mutations and the distribution of their effects on fitness change across environments (Martin and Lenormand 2006b; Wang *et al.* 2009; Agrawal and Whitlock 2010; Wang *et al.* 2014). In particular, the fraction of beneficial mutations was found to increase in stressful environments (Remold and Lenski 2001, 2004). Standing genetic variation for quantitative traits (notably fitness components), also frequently depends on the environment (Hoffmann and Merilä 1999; Sgrò and Hoffmann 2004; Charmantier and Garant 2005). Finally, the initial frequency of preexisting variants able to rescue the population from extinction in a stressful environment may depend on their selective cost in the past environment. In light of this empirical evidence, it seems clear that progress towards understanding and predicting ER across stress levels requires addressing, in a quantitative way, the joint effect of stress on the demography and genetic variation in fitness of a population exposed to stressful conditions. This is our goal in the present article.

To do so, we develop a model that is a hybrid between two modeling traditions in ER theory, summarized by Alexander *et al.* (2014): discrete genetic models, and quantitative genetic models. Discrete genetic models assume a narrow genetic basis for adaptation (and ER), whereby a single beneficial mutation can rescue an otherwise monomorphic population (Orr and Unckless 2008; Martin *et al.* 2013; Uecker *et al.* 2014; Orr and Unckless 2014; Uecker and Hermisson 2016). This approach was initially proposed for ER by Gomulkiewicz and Holt (1995), and later extended to account for (i) evolutionary and demographic stochasticity (e.g. Orr and Unckless 2008), and (ii) variation in the selection coefficients of mutations that may cause rescue, with an arbitrary distribution of fitness effects (Martin *et al.* 2013). However, such models do not predict how the distribution of fitness effects of mutations vary along gradients of stress level. For this reason, they make it difficult to jointly address the two fundamental components of stress mentioned above. On the contrary, quantitative genetics models of ER inherently address the influence of stress on the rate of adaptation by assuming that adaptation (and ER) is caused by evolution of a quantitative trait whose optimum changes with the environment (Lynch *et al.* 1991; Burger and Lynch 1995; Gomulkiewicz and Holt 1995). In these models, both the rate of population decline and the rate of adaptation under stress depend on the distance between the phenotypic optima in the past and present environments. However, analytical predictions are derived assuming a broad, polygenic basis for adaptation with a stable genetic variance of the quantitative trait. The population genetic processes underlying adaptation are not explicitly modelled, and the stochasticity involved in fixation and establishment of mutations neglected. These complications are only explored by simulations (e.g. Gomulkiewicz *et al.* 2010).

In order to take the best of both approaches, we rely on Fisher’s (1930) Geometrical Model (hereafter “FGM”). Fitness variation in the FGM is assumed to emerge from variation in multiple putative phenotypic traits undergoing stabilizing selection that depends on the environment. This model is analytically tractable, while retaining various aspects of realism (reviewed in Tenaillon 2014). In particular, it accurately predicts how fitness effects of mutations change across environments (Martin and Lenormand 2006b; Hietpas *et al.* 2013; Harmand *et al.* 2017) or genetic backgrounds (Martin *et al.* 2007; MacLean *et al.* 2010; Trindade *et al.* 2012). The FGM naturally relates environmental stress to (i) the rate of population decline, (ii) the rate and effect of rescue mutants, and (iii) their potential costs in the past environment. Here, we combine this FGM with population dynamic approaches that account for demographic and evolutionary stochasticity (Martin *et al.* 2013), in a regime where selection is strong relative to the rate of mutation. We consider rescue in asexual populations, stemming either from *de novo* mutations or standing genetic variance. Interestingly, we show that all effects of stress on demography and on the distribution of the fitness effects of mutations can be summarized into a single composite measure of effective stress level. Evolutionary rescue shifts abruptly from very likely to very unlikely over a narrow window of effective stress level, which can be predicted from empirically measurable quantities.

## METHODS

We here detail the ecological (environmental), genetic, and demographic assumptions of the model, and the approximations used for its mathematical analysis.

### Abrupt environmental shift

We define two environments: (1) a non-stressful one, denoted as “previous environment”, in which the population has a positive mean growth rate, and a large enough population size that demographic stochasticity can be ignored; and (2) a stressful one, denoted “new environment”, in which the population initially has a negative mean growth rate, and the population size is subject to demographic stochasticity. Conditions shift abruptly from the previous to the new environment at ***t*** = **0**, at which time the population size is *N*_0_.

### Eco-evolutionary dynamics

Extinction or rescue ultimately depends on details of the stochastic population dynamics of each genotype. These are assumed to be mutually independent (no density or frequency-dependence, see Chevin 2011), and sufficiently ‘smooth’ (moderate growth or decay) that they can be approximated by a Feller diffusion (Feller 1951), following Martin *et al.* (2013). This approximation reduces all the complexity of the life cycle into two key parameters for each genotype ***i***: its expected growth rate ***r_i_*** (our fitness here), and its variance in reproductive output ***σ*_*i*_**. Our simulations below are performed for discrete generations with Poisson offspring distributions. In this case, ***σ_i_* = 1 + *r*_*i*_ ≈ 1** for any genotype, as long as their growth rate is not too large (***r*_*i*_** ≪ 1 per-generation, see **Appendix section I subsection 1 and 2** and Martin *et al.* (2013)). Note that the approximation extends to various other forms of reproduction (see Martin *et al.* 2013).

To cause a rescue, a resistant mutant (***r*_*i*_ > 0**) must establish, by avoiding extinction when rare. The probability that this happens, for a lineage with growth rate ***r*_*i*_ > 0** starting from a single copy is **1 − *e^−2r_i_^*** (still assuming ***r*_*i*_ ≪ *σ*_*i*_**, with ***σ*_*i*_, ≈ 1** in the example used in simulations). The number of individuals from which such mutations can arise declines in time, and we ignore stochasticity in these decay dynamics. This is accurate as long as the population has large initial size, of order ***N*_0_ ≫ 1** (Martin *et al.* 2013).

Finally, we assume that mutation rates per capita per unit time are constant over time. This is exact in models with discrete generations. In continuous-time models, where mutations occur during birth events, mutation rates vary between genotypes with different birth rates, and over time as these genotypes change in frequency. However, the constant mutation rate model can still be approximately valid (see Martin *et al.* 2013).

### ER from standing variance versus *de novo* mutation

At the onset of stress **(*t* = 0)**, the population either consists of a single ancestral clone, or is polymorphic at mutation-selection balance in the previous environment. In the first case, we must derive the distribution of fitness effects, in the new environment, of mutants arising from the ancestral clone. In the second case, we must also describe the potential rescue variants already present in the previous environment.

### Mutations under Fisher’s geometrical model (FGM)

We assume that the expected growth rate of a given genotypic class [its Malthusian fitness, or log-multiplicative fitness in discrete-time models), is a quadratic function of its phenotype for ***n*** quantitative (continuous) traits. Denoting as **z** ∈ ℝ^*n*^ the vector of breeding values (heritable components) for all traits, and as **o** the single optimum phenotype with maximal growth *r_max_*, the expected growth rate is

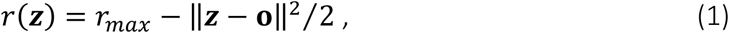

while the stochastic variance in reproductive success is assumed constant across genotypes.

The key assumption of our model is that the optimum depends on the environment. Without loss of generality, we set the phenotypic origin at the optimum in the new environment, in which **o = 0**. In the previous environment, the optimum coincides with the mean phenotype of the ancestral population (‘A’): **o = z_*A*_ = 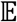(z)**, which implies that the ancestral population was well-adapted in its original environment. The fitness of the mean ancestral phenotype **z_*A*_** in the new environment is thus 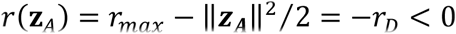, where *r_D_* is its rate of decay, and the phenotypic magnitude of the stress-induced shift of the optimum phenotype [from ***o = z_A_*** to **o = 0**) is 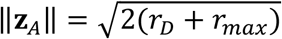.

Mutations occur as a Poisson process with rate *U* per unit time per capita, constant over time and across genotypes, but potentially variable across environments. Each mutation adds a random perturbation **dz** to the phenotype, drawn from an unbiased and isotropic multivariate Gaussian distribution **dz ~ *N*(0, *λ* I_*n*_)**, where **I_*n*_** is the identity matrix in *n* dimensions and *λ* is a scale parameter. Note that, since traits are not our main interest here, we choose to measure mutation effects on them in units that directly relate to their fitness effects. Therefore, *λ* can be understood as the variance of mutational effects on traits, standardized by the strength of selection (see **Appendix section II subsection 1** for more details).

Note that mutation effects are additive on *phenotypes* (no epistasis), but not on *fitness*, because ***r*(z)** is nonlinear (Martin *et al.* 2007).

**Fig.1** illustrates the rescue process in the FGM. At the onset of stress (*t* = 0), the optimum shifts abruptly to a new position, such that the mean growth rate becomes negative with **−*r_D_* < 0** (Fig.1.C). Meanwhile, the population size starts to drop from an initial value ***N*_0_** (Fig.1.C), facing extinction in the absence of evolution. However, one or several mutants or pre-existing variants may be close enough to the new optimum to have a positive growth rate (“resistant genotypes”, **Fig.1.A, 1.B**). These may then establish, and ultimately rescue the population (“rescue genotypes” **Fig.1A, 1.B**).

**Figure 1:**
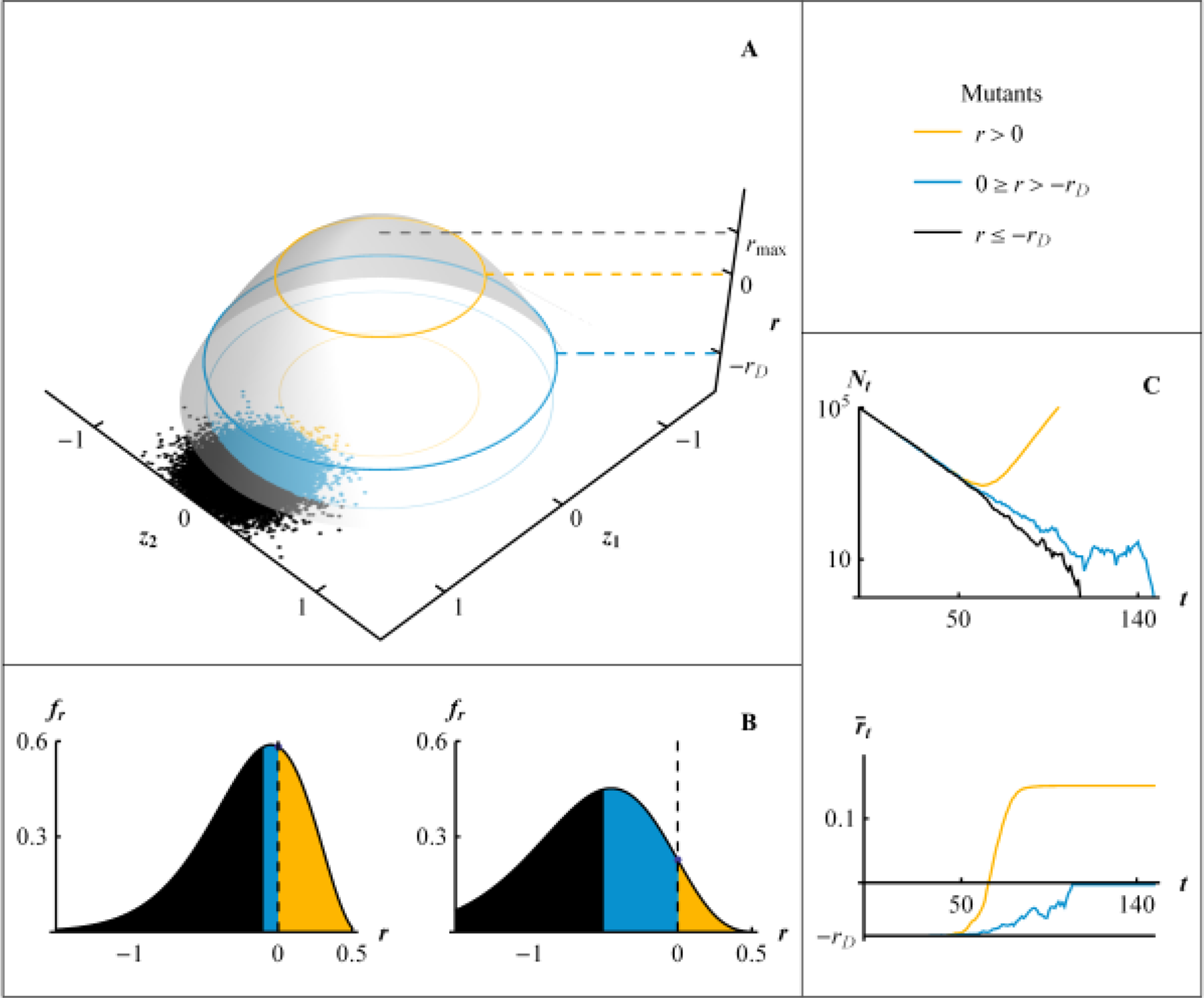
Evolutionary rescue in Fisher’s geometric model. In all panels, black refers to deleterious and neutral mutations,(−*r*_*D*_ ≥ *r*) blue to beneficial but not resistant mutations (−*r*_*D*_ < r ≤ 0) and orange to resistant mutations (*r* > 0), around the dominant genotype of the ancestral population with phenotype *z*_*A*_ ≠ 0. (A) Fitness landscape (FGM) with growth rate *r* (*z*-axis) determined by two phenotypic traits *z*_1_ and *z*_2_. Dots represent the distribution of random mutant phenotypes around the dominant genotype of the ancestral population. The growth rate of this dominant genotype, in the stressful environment, is −*r*_*D*_, and *r*_*max*_ is the maximal fitness at the phenotypic optimum. (B) Distribution of growth rates among random mutants arising from the dominant genotype (distribution of mutation effects on fitness) for two decay rates *r_D_* = 0.1 (left) and *r_D_* = 0.5 (right). (C) Dynamics of the population size *N*_*t*_ and mean fitness 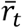 of a population starting from a clone at −*r_D_* = −0.083 at size *N*_0_ = 10^5^. The black line represents the case without fixation of a beneficial mutation, the blue line the case with extinction in spite of the fixation of a beneficial, but non-resistant, mutation, and the orange line the case of a rescue. Parameters for the simulations are *r_max_* = 1.5, *U* = 2 * 10^−5^, *n* = 4 and *λ* = 5 * 10^−3^.

Within the context of the FGM, increasing stress level may have different effects, also discussed in Harmand *et al.* (2017). First, stronger stress may cause a larger shift in the position of the optimum phenotype, resulting in a larger initial drop in fitness (higher *r_D_*), as assumed in most models of adaptation to a changing environment (Kopp and Matuszewski 2014). In addition, the maximal possible fitness *r_max_* may also be lower in the new than in the previous environment (reduced environmental quality). Moreover, the mutational parameters (*U* and *λ*) may change with stress, causing shifts in evolvability. Note that a change in *λ* may reflect a change in the phenotypic effects of mutations, of the strength of stabilizing selection, or both. For instance, higher stress may release cryptic genetic variance on underlying phenotypic traits (Scharloo 1991; Hermisson and Wagner 2004), or cause increased mutation rates via SOS responses in bacteria (Foster 2007). Finally, although less easy to conceptualize, some environments may change the effective dimensionality of the landscape. However, in the present paper, we only consider such changes in dimensionality in the context of rescue from *de novo* mutations (where it can readily be handled by studying the effect of the parameter *n*).

As we will see, all our results can be expressed in terms of five parameters (*N*_0_*U*, *r_D_, r_max_, λ,n*. Table 1 summarizes all notations in the article.

**Table 1:**
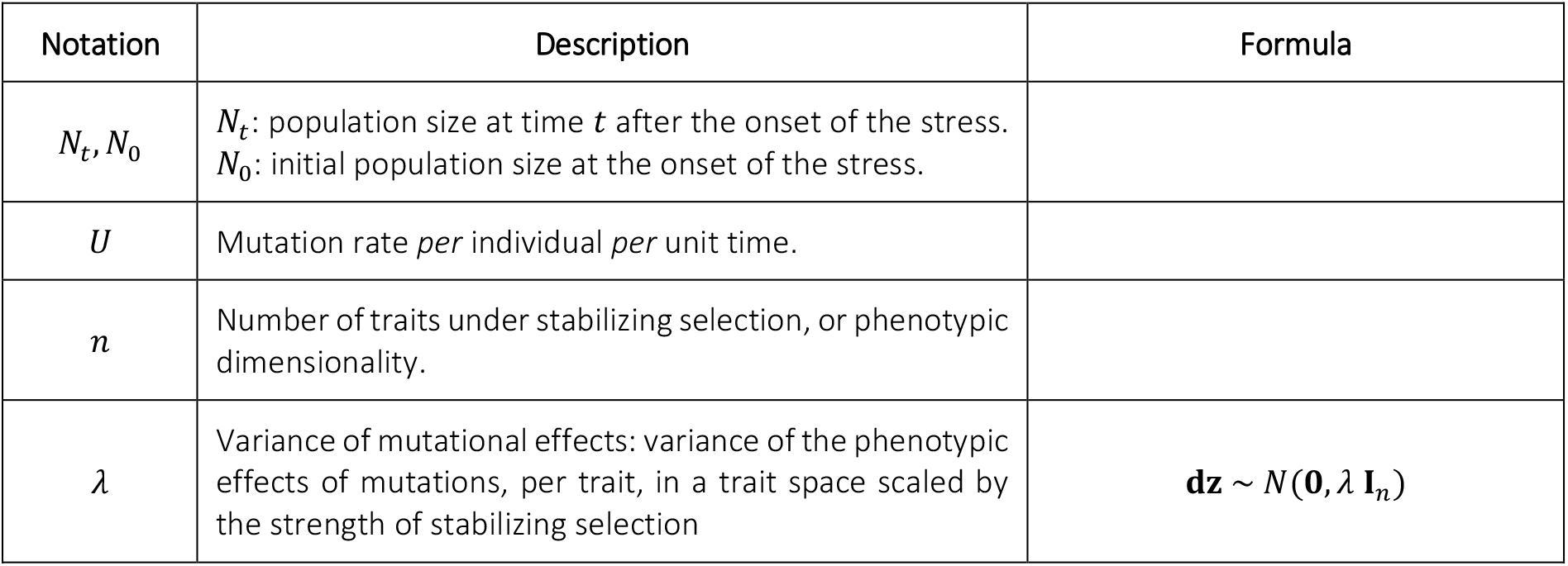
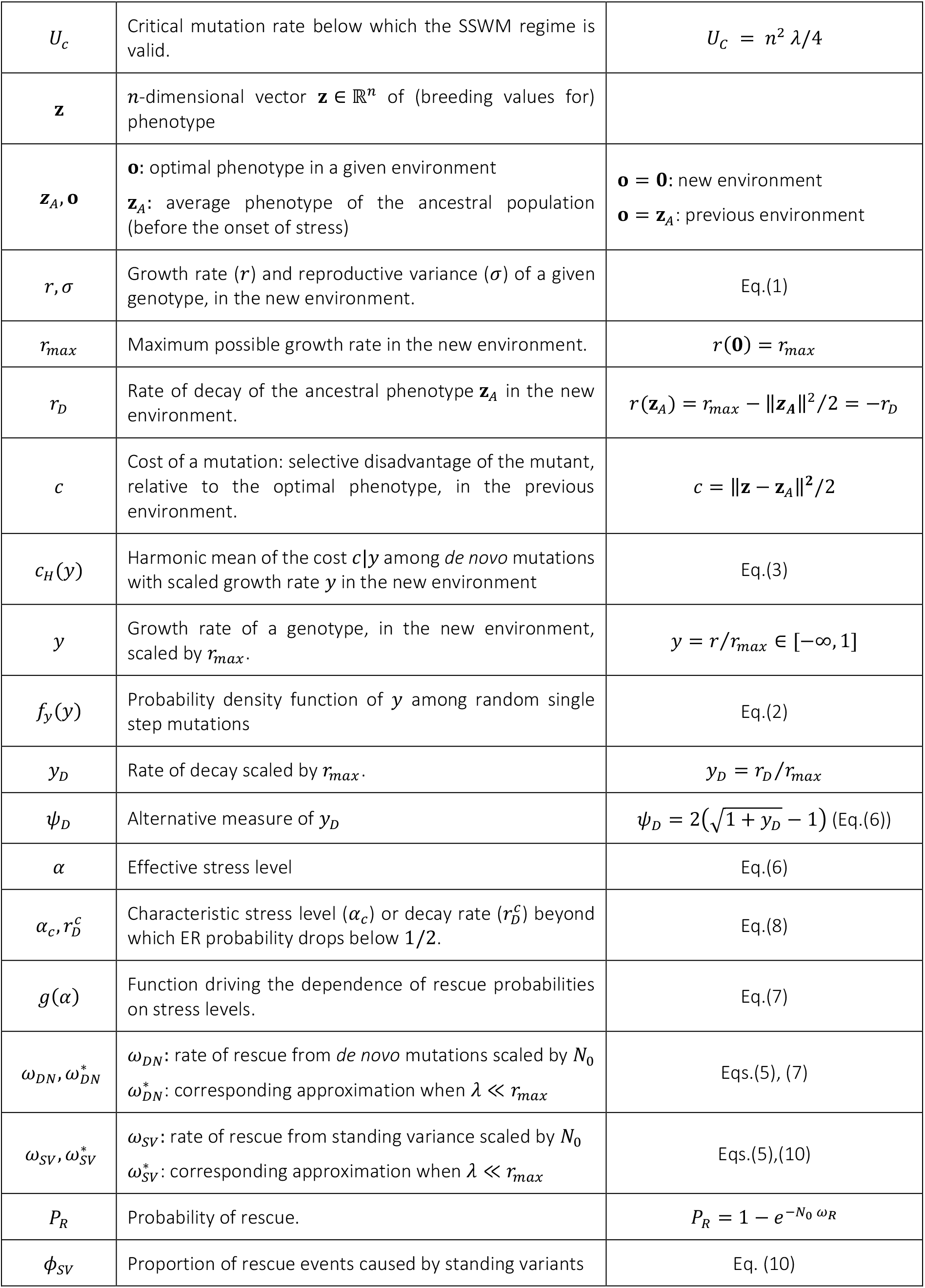
Notations

### Strong selection and weak mutation (SSWM) regime

The FGM described in the previous section, produces epistasis for fitness between different mutations which makes the problem highly intractable in general. To make analytical progress, we assume a regime of strong selection and weak mutation (SSWM, Gillespie 1983), which allows neglecting multiple mutants and epistasis. This regime arises when mutation rates are small relative to their typical fitness effect (as detailed below). In our context, this assumption implies that most rescue variants (pre-existing or *de novo*) are only one mutational step away from the ancestral genotype, allowing for two key simplifications. First, with a purely clonal ancestral populations, we can ignore ER by genotypes that have accumulated multiple *de novo* mutations. Second, in populations initially at mutation-selection balance, we can consider that all mutations arise from a single dominant genotype, optimal in the previous environment. Indeed, in the SSWM regime at mutation-selection balance, most segregating phenotypes remain within a narrow neighborhood of the optimum (relative to the magnitude of mutation effects on traits), so the mutation-selection balance is well-approximated by assuming that all mutations originate from the optimum phenotype. This is essentially the House-of-Cards approximation (Turelli 1984) extended to the FGM of arbitrary dimensionality (Martin and Roques 2016).

Overall, the SSWM assumption implies that the evolutionary aspects of ER are entirely determined by a single joint distribution of fitness, in the previous and new environment. This distribution corresponds to that of mutants arising from the optimal genotype of the previous environment. We thus apply the results of Martin *et al.* (2013), to this particular distribution.

Note that the SSWM approximations used in this article should apply even when multiple single-step mutants co-segregate (generating “soft selective sweeps”, as detailed in Wilson *et al.* 2017). Indeed, the probability of ER, as computed for example in Orr and Unckless (2008) or Martin *et al.* (2013) and used here, is one minus the probability that no *single* mutant arises that ultimately causes ER. This means that we ignore ER requiring multiple mutational steps, but allow several single-step rescue mutations to co-segregate. Consistently, our simulations did not show any particular deviation from the theory at very mild stress, where such co-segregation of several single-step mutants is expected.

### Maximal mutation rate for the SSWM regime

We conjecture that the SSWM approximation should be accurate below some threshold mutation rate *U*_*c*_, i.e. whenever *U* < *U_c_* = *n*^2^ *λ*/4.

Indeed, Martin and Roques (2016) found that as long as *U ≤ U*_*c*_, the fitness distribution at mutation-selection balance corresponds exactly to that expected under the House of cards approximation (with a dominant optimal genotype plus its deleterious mutants). Whether this same condition is sufficient for most rescue events to stem from single-step mutations is not justified theoretically, and was simply tested by extensive stochastic simulations. **Supplementary Fig. 1** further explores the range of validity of this approximation. It shows, in a rescued population, the proportion of wild-type, single mutant, double mutant, and so on, as a function of the mutation rate.

### Distribution of single-step mutation fitness effects in the new environment

Let *s* be the selection coefficient (difference in growth rate), in the new environment, of a random mutant (with phenotype **z**) relative to its ancestor (with phenotype *z_A_)*. The distribution of *s* = *r*(**z**) − *r*(**z**_A_) = *r* + *r*_*D*_ among random mutants has a known exact form in the isotropic FGM (Martin 2014; Martin and Lenormand 2015), from which the distribution of growth rates (*r* = *s* – *r*_*D*_) in the new environment is readily obtained. It proves simpler and sufficient (see **Appendix section II subsection 2**) to consider the scaled (and unitless) growth rate *y* = *r/r_max_* Σ [−∞, 1], such that *y_D_* = *r_D_/r_max_* ∈[0, +∞] is the decay rate of the ancestor scaled to the maximum possible growth rate. The scaled growth rates *y* = *r/r_max_* have the following probability density function:

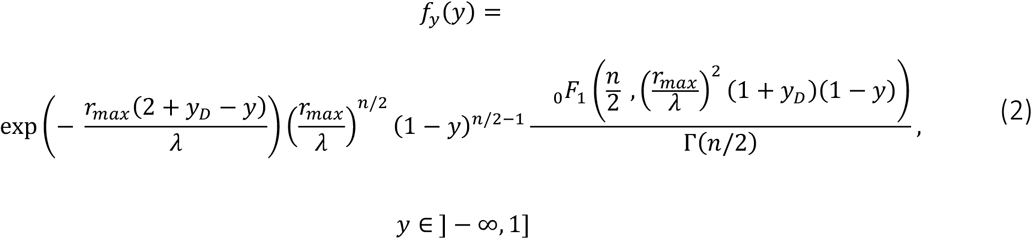

where _0_*F*_1_(.,.) is the confluent hypergeometric function and Γ(*z*) is the gamma function. In the SSWM regime, this probability density function approximately describes *de novo* mutations produced after the onset of stress by the whole population, be it initially clonal or at mutation-selection balance.

### Fitness cost of single-step pre-existing mutants in the previous environment

Consider the subset of random mutations, among those that arise from the dominant genotype of the ancestral population, that have a scaled growth rate within the infinitesimal class [*y,y* + *dy*] in the new environment. We introduce the conditional random variable *c*|*y*, which is the cost, in the previous environment, of a random mutant within this subset (thus, conditional on *y*). This cost is equal to the negative of the selection coefficient of the mutation relative to the dominant genotype with phenotype ***z***_*A*_. More precisely, the cost of a mutant with phenotype **z** is 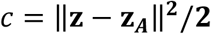 (using eq.(1), with **o = z_*A*_** for the previous environment). Note that, because the mutation-selection balance in the previous environment is fully characterized by relative fitnesses, which do not depend on the maximal growth rate in this environment, the latter may differ from *r_max_* without impacting the distribution of the costs *c*|*y* and our results. Importing results from Martin *et al.* (2013) for the SSWM regime, the total number of preexisting variants within the class [*y,y* + *dy*] is Poisson distributed with mean ***N*_0_ *U f_y_(y)/c_H_(y)dy***, where ***c_H_(y)*** = 1/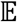_*c*_(**1/*c*|*y***) is the harmonic mean of the cost c|y among mutations with effect *y* in the new environment. This conditional harmonic mean depends on the joint distribution of mutation effects on fitness (*c, y*) across two environments in the FGM (given in Martin and Lenormand 2015). In our context, the dominant genotype of the ancestral population is optimal in the previous environment and far from the optimum in the new environment. In this case, using Eq.(9) in Martin and Lenormand (2015), the resulting conditional harmonic mean *c_H_(y*) takes a tractable form (see Eq.(A6) for *n* ≥ 2 and (A8) for ***n* = 1** in **Appendix section II subsection 4 and 5**):

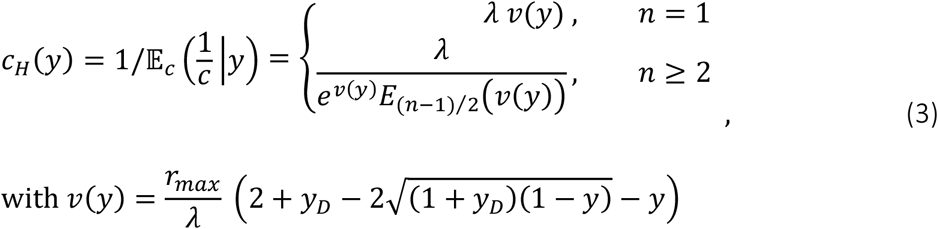

where 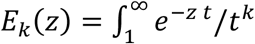 is the exponential integral function. In most of the article we focus on the case *n* ≥ 2, when considering ER from standing variance. The distributions of mutation effects on fitness in both the previous (Eq.(3)) and the new environment (Eq.(2)) can then be integrated to yield the probability of ER, as we show next.

#### General expression and assumptions for the rescue probability

Extinction occurs when no resistant mutation manages to establish (i.e. to avoid stochastic loss). For compactness, we define a rate of rescue *ω* per individual present at the onset of stress (i.e., scaled by *N*_0_), such that, following Martin *et al.* (2013), ER probabilities take the general form (similar to that in e.g. Orr and Unckless 2008):

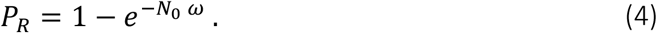

The rate of rescue from *de novo* mutations alone is *ω_*DN*_* (‘*DN*’ for *de novo*), while that from pre-existing variance alone is *ω*_*sv*_ (‘*SV*’ for standing variants). For a purely clonal population, the rate of rescue is *ω* = *ω_DN_*, while for a population initially at mutation-selection balance, it is *ω = ω_DN_ + ω_sv_* in the SSWM regime assumed here (Martin *et al.* 2013). Applied to the context of the FGM using Eqs.(2) and (3), the rates *ω_DN_* and *ω_sv_* are given by (see **Appendix** Eq.(A5) and (A7)):

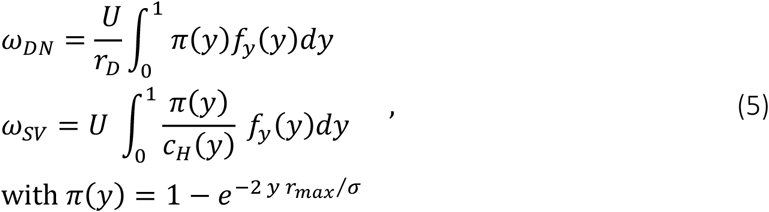

where *c_H_(y*), and *E*_(*n*-1)/2_(.) are defined in Eq.(3) and *π*(*y*) is the probability of establishment of a resistant genotype with scaled growth rate *y* > 0 in the new environment. *ω_DN_* in Eq.(5) is simply the average establishment probability of *de novo* resistant mutants times the genomic mutation rate, divided by the rate of decay. In previous ER models (e.g. Orr and Unckless 2008; Martin *et al.* 2013), which we denote “context-independent”, the probability of rescue takes the exact same form as Eq.(4). The expressions for the rates of rescue per capita also take a form similar to Eq.(5): for *de novo* mutations, *ω_DN_ = U q_R_ / r_D_*, and for standing variance, *ω_sv_* = *U q_R_ / c_H_*, where *q_R_* is the proportion of rescuers among random mutations (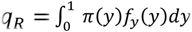 in Eq.(5)) and *c_H_* is again the harmonic mean of the cost of rescue mutations. The important difference is that in previous models, *q_R_* and *c_H_* do not depend on *r_D_*, while the corresponding quantities in Eq.(5) do depend on the rate of decay, through its effect on *f_y_(y)* and *c_H_*(y).

The linearity of ER rates with the mutation rate *U* (*ω* ∝ *U*) arises here because of the SSWM regime, where multiple mutations are ignored: it might not hold at higher mutation rates (when *U* > *U_c_*). As such, Eq.(5) makes no further assumption than the SSWM regime (*U < U_c_*); it can easily be evaluated numerically to provide a general testable theory for rescue probabilities across stress levels, in the FGM. Yet, in order to gain more quantitative/intuitive insight into the effects of stress, we study approximate closed forms for the rates in Eq.(5).

#### Small mutational effects approximation (SME)

Although selection is assumed to be strong relative to mutation (*U < U_c_*, SSWM regime), it is still fairly realistic to assume that mutation effects on traits (and thus fitness) are weak relative to the maximal growth rate in the new environment, namely that *λ* ≪ *r_max_*. Taking a limit where *λ/r_max_* → 0, simpler expressions for Eq. (5) are derived in the **Appendix section III**.

With this approximation, single-step resistance mutations are still rare and of large phenotypic effect, in that they pertain to the tail of the mutant phenotype distribution. However, even resistance mutations typically remain far from the optimum in the new environment, so that their scaled growth rate is small: *y = r/r_max_* ≪ 1. Overall, mutation effects must fall within the range: 4*U/n*^2^ < *λ ≪ r_max_* for both the SSWM and the SME (small mutational effects) approximation to apply (see **Appendix section** III). In the appendix, we study the convergence, as *λ/r_max_* decreases, of the results from Eq.(5) to their asymptotic limit (**Supplementary Figs.3 and 4**).

#### Stochastic simulations of a discrete-time model

We checked the robustness of our assumptions and approximations using stochastic simulations, where we tracked the population size and genetic composition of a population across discrete, non-overlapping generations. The size *N_t+1_* of population at generation *t + 1* was drawn as a Poisson number 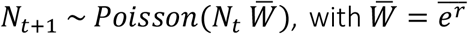 the mean multiplicative fitness (*W = e^r^*) and *N_t_* the population size, in the previous generation. The genotypes forming this new generation were then sampled with replacement from the previous one with weight *W_i_* = *e^r_i_^*. This is faster and exactly equivalent to drawing independent Poisson reproductive outputs for each individual, or genotype. Because of the underlying assumptions of the simulations, the corresponding analytical approximation for the stochastic reproductive variance in Eq.(1) is σ_i_ = σ ≈ *1* (assuming small growth rates *r_i_* ≪ 1). Mutations occurred according to a Poisson process, with a constant rate *U* per capita per generation. Mutation phenotypic effects were drawn from a multivariate normal distribution ***N*(0, *λ* I_*n*_)**, with multiple mutants having additive effects on phenotype, and their fitness computed according to the FGM (Eq.(1)).

Rescue probability was estimated by running 1000 replicate simulations until either extinction or rescue occurred. A population was considered rescued when it reached a population size *N_t_* and mean growth rate 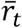 such that its ultimate extinction probability, if it were monomorphic, would lie bellow 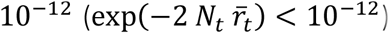. This is a conservative criterion: once 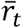 has become positive, we expect it to remain so, yielding further increases in population size and thus further decreasing the probability of future extinction. We checked on a subset of simulations that the above procedure gave the same rescue probabilities as obtained in simulations performed until the population rebounded back to its (large) initial size *N_0_*.

For rescue from populations at mutation-selection balance, 8 replicate initial equilibrium populations were generated, each by starting from an optimal clone and running the same algorithm with fixed population size (*N_t_* = 10^6^) until the mean growth rate had visually stabilized to a fixed value (close to its theoretical equilibrium value 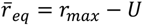 (for *U < U_c_*) for more than 1000 generations. Then the optimum was shifted by 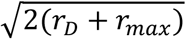 phenotypic units, and 1000 replicate ER simulations were performed (same algorithm as for *de novo* rescue), from each of the 8 replicate equilibrium populations.

All simulations and mathematical derivations were performed in *MATHEMATICA* v. 9.0 (Wolfram Research 2012).

## RESULTS

The ER rates in Eq.(5) are analytical but only implicit functions of the model parameters. In a small mutational effects (SME) limit, they take simpler closed form (indicated by a ’;*’). As we will see below, these simpler forms mostly depend on the following two compound variables, which summarize the various effects of stress on the fitness landscape:

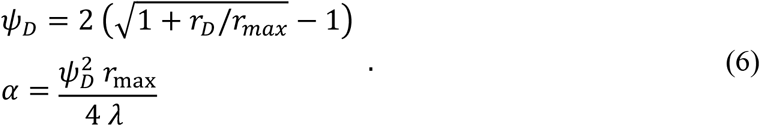

Both and *ψ_D_* and α increase with the decay rate *r*_D_, decrease with increasing peak height *r*_max_, and are independent of *n*. The parameter *α* further increases with decreasing variance of mutational effects *λ*. We can already see how *α* qualitatively reflects an “effective stress level”: stress is harder to cope with if decay rate is larger, the maximum growth rate is lower, and mutation effects are smaller.

### Rescue from *de novo* mutations

Under the SME approximation and in the SSWM regime (4 *U/n^2^ < λ ≪ r*_max_) the rate of *de novo* rescue (Eq.(5)), converges to (Eq. (A12) in the Appendix):

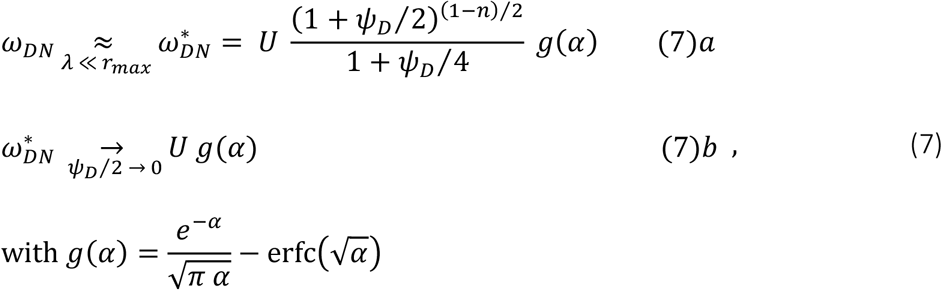

where erfc(.) is the complementary error function. Eq.(7)*b*. gives the approximate closed form of Eq.(7)*a*. for mild stress (*ψ_D_*/2 → 0). Note that this approximation converges faster (with decreasing ψ_D_) with fewer dimensions, due to the faster vanishing of the factor (1 + *ψ_D_/2)^(1-n)/2^* (in the limit *n* = 1 it vanishes for all ψ_D_). We now discuss the biological implications of these expressions.

### Effect of FGM parameters on rescue

The partial derivatives of *ω*_DN_* in Eq.(7) with respect to the FGM parameters (*r_D_, r_max_, λ, n*) quantify the sensitivities of ER probability to each of them (Appendix section III subsection 4). First, note that *g*(.) is a strictly decreasing function of *α*. When *n > 1* and with mild stress (*ψ_D_* ≪ 2), Eq.(7)b. applies and *ω_DN_* ≈ *U g(α)*. ER then becomes less likely with a higher decay rate *r*_D_, a lower peak *r_max_* and a smaller variance of mutational effects *λ*, and is independent of dimensionality *n*. For stronger stress levels, Eq.(7)*a*. applies: these qualitative dependencies to the parameters still hold, except that ER probability now decreases with increasing dimensionality.

### Sharp drop of ER probability with stress levels

Fig.2 shows the agreement between simulations (stochastic discrete-time demographic model, see Methods) and the analytical expressions in Eqs.(5) and (7), over a wide range of stress levels (quantified by *r*_D_), and for two values of *r_max_* and *U* (Supplementary Fig.3 further explores the range of validity of the approximation). Interestingly, ER probability drops sharply with stress levels (with decay rate *r_D_* here), which is well captured by the term *g*(*α*) alone (Eq.(7)*b*., dashed red lines in **Fig.2**). This drop is much more pronounced than in a context-independent model (gray lines in **Fig.2**), where stress does not affect the distribution of mutation effects. The difference between context-independent models and the FGM is that, in the latter, increased stress implies both faster decay (as in the former), and fewer and weaker resistance mutations. In the FGM, these effects on the properties of rescue mutations are the main drivers of ER probabilities across stress levels.

**Figure 3:**
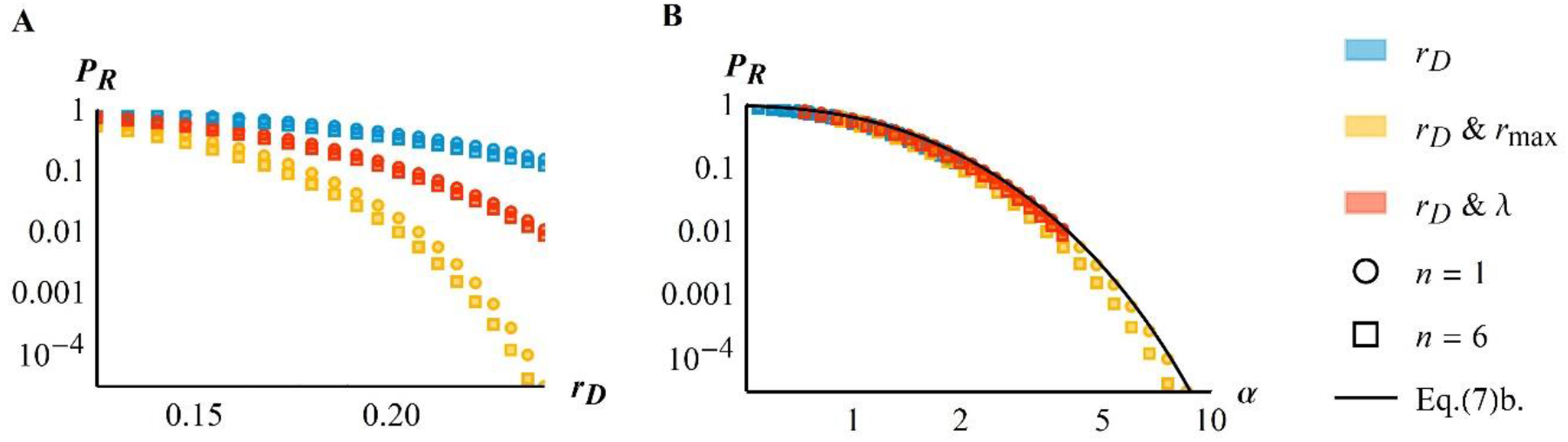
The effective stress level. Rescue probability for clonal populations versus initial decay rate r_D_ (A) or the effective stress level α (B). In both panels the axes are in logarithmic scale, symbols show Eq.(5) and colors refer to different effects of increased stress level: blue symbols show only r_D_ increasing (with r_max_ = 1.5 and λ = 0.005), orange symbols r_D_ increasing and r_max_ decreasing linearly with r_D_ (according to r_max_=1.5−5 r_D_, with λ = 0.005) and red symbols r_D_ increasing and λ decreasing linearly with r_D_ (according to λ = 0.005 − 10^−2^ r_D_, with r_max_ = 1.5). In each case, the results for both *n* = 1 (circles) and *n* = 6 (squares) are shown. The black plain line on the right panel gives the result from Eq.(7)b.: a single composite measure of stress (α) approximately captures the impact of stress-induced variations in the various parameters (r_D_,r_max_,λ,n). Other parameters are *N*_0_ = 10^6^ and *U* = 2 * 10^−5^.

**Figure 2:**
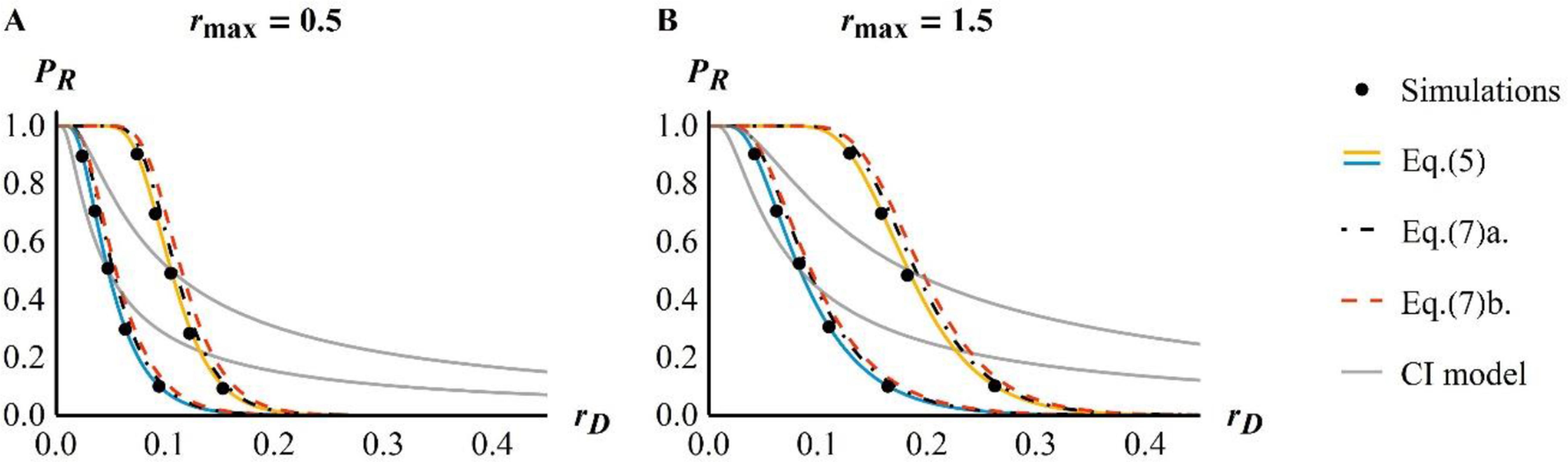
Rescue probability from *de novo* mutations. The ER probability as a function of stress levels, expressed as the initial mean decay rate of the population, is given for various values of the mutation rate *U* = 10^−3^ *U*_c_ (blue) or *U* = 10^−2^ *U*_c_. (orange) and the maximal fitness reachable in new the environment r_max_ = 0.5 (A) or 1.5 (B). Dots give the results from simulations and solid lines (blue and orange) show the corresponding theory computed numerically (Eq.(5)). The black dot-dashed (respectively red dashed) lines give the corresponding analytical approximations Eq.(7)a. (respectively Eq.(7)*b*.). The gray lines correspond to an equivalent theory without the FGM (named context-independent model as described in the Method section, “CI”) modified from Orr and Unckless (2008). This last model was computed using a fixed proportion of resistant mutations equal to the one in Eq.(5) for a rescue probability of 0.5 (which explains why the two curves cross exactly at P_R_ = 0.5). Other parameters are n = 4, *N*_0_ = 10^5^, λ = 0.005.

### Composite parameter *α* describing an effective stress level

The results in Eqs.(6)-(7) suggest that a single composite parameter (*α* in Eq.(6)) can capture the various ways in which stress may alter the parameters of the fitness landscape, that is, fitness peak height *r*_max_, variance of mutational effects *λ*, or distance to the optimum *r*_D_. We denote this parameter the “effective stress level”. In **Fig.2**, considering the effect of stress only via *α* (Eq.(7)b.) is equally accurate as using the more complex Eq.(7)a., or the numerical computation from Eq.(5).

This simplification is further illustrated in **Fig.3**, where we use exact numerical computations from Eq. (5) to explore different possible effects of stress. Regardless of whether stress affects only the maladaptation of the ancestral clone (***r_D_***, blue symbols), or also the quality of the environment (joint change in ***r_D_*** and ***r_max_***, orange symbols) or the variance of mutational effects (joint change in ***r_D_*** and *λ*, red symbols), its effect on the rescue probability is accurately predicted by *α* (**Fig.3B**, black line). As predicted by Eq.(7)b., the relationship between rescue probability and *α* is approximately independent of dimensionality (compare circles *n = 1* and squares *n = 6* in **Fig.3B**). We also note that Eq.(7)b. slightly overestimates the ‘exact’ numerical computations of the ER probability from Eq.(5), so it provides a conservative bound when considering the control of resistant pathogens.

### Characteristic stress level

Fig.2 shows that ER drops from highly likely to highly unlikely around a “characteristic stress level”, which we can be characterized analytically (as detailed in the **Appendix section IV subsection 1**). Consider the set of values of parameters (***N_0_, U, r_max_, r_D_,λ, n***) for which the rescue probability is of given value **p ∈ [0,1]**. From Eq.(7)b., this corresponds to the set (*r_max_, r_D_, λ*) for which *α = g^−1^(−log(1 — p*) /*N_0_U*). Using the approximation 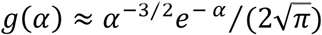 (from Eq.(7) with *a* α ≫ 1), the corresponding *α* can be derived explicitly (Eq.(A17)). In particular for p = 1/2, the characteristic stress level *α_c_* at which the ER probability is 1/2 is (Eq. (A19) in the **Appendix**):

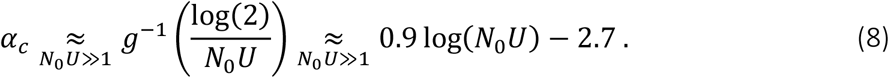

Under the conditions of the SME approximation (detailed in Methods), Eq.(8) applies for large *N_0_U* (approximately when *N_0_U ≥* 5.10^4^), a necessary condition for this equation to be selfconsistent (detailed in **Appendix section IV subsection 2**).

The characteristic stress level *α_c_* that a population can typically withstand increases only log-linearly with population size and mutation rate. Consider the characteristic decay rate 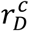 for which the rescue probability is *P_R_ = 1/2*, i.e. the decay rate that populations can overcome half of the times. From Eq.(8) with 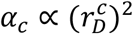 (Eq.(6)), this decay rate is 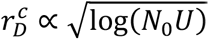 for large *N_0_U*. For comparison, we would have 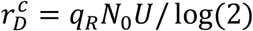 which is linear in *N_0_U*, in a context-independent model where the proportion *q_R_* of random mutations causing a rescue is independent of *r_D_*. The difference in the effect of *N_0_U* on rescue probability thus stems from the strong non-linearity (i.e. sharp drop) of rescue probability with stress level (decay rate) under the FGM. In the FGM, overcoming a given environmental harshness requires much more mutational input than in a context-independent model.

### Characteristic stress window

It is also important to predict how sharply the ER probability drops around the characteristic stress level. This drop can be characterized by a “characteristic stress window” of *α* over which the ER probability drops from 75% to 25%. The width *Δα* of this window can be scaled by the value of the characteristic stress level *α*_*c*_, to get a scale-free measure of its steepness (i.e., how sharp the drop in ER probability is, relative to the stress level around which it occurs). This gives (from Eq.(A20) in the **Appendix**):

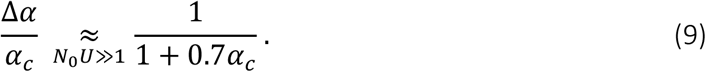

The width Δ*α* increases with increasing *N*_0_*U* until it saturates (at ~ **1.5**) (see **Appendix section IV subsection 3** for further details). However, when scaled by the center of the window (*α_c_*), the width of this scaled characteristic stress window drops below 1 as *α_c_* increases (and hence with increasing *N_0_U*, from Eq.(8)). This result shows formally that the drop in ER probability with increasing stress *α* gets proportionally sharper (relative to the position where it occurs) as *N_0_U* increases, but this is entirely driven at large *N_0_U* by shifts of a window with constant absolute width. This is illustrated in **Fig.4**, which also shows the accuracy of Eq.(9) compared to ‘exact’ numerical computations from Eq.(5) (as expected, the exact result deviates from Eq.(9) for smaller *N_0_U)*.

**Figure 4:**
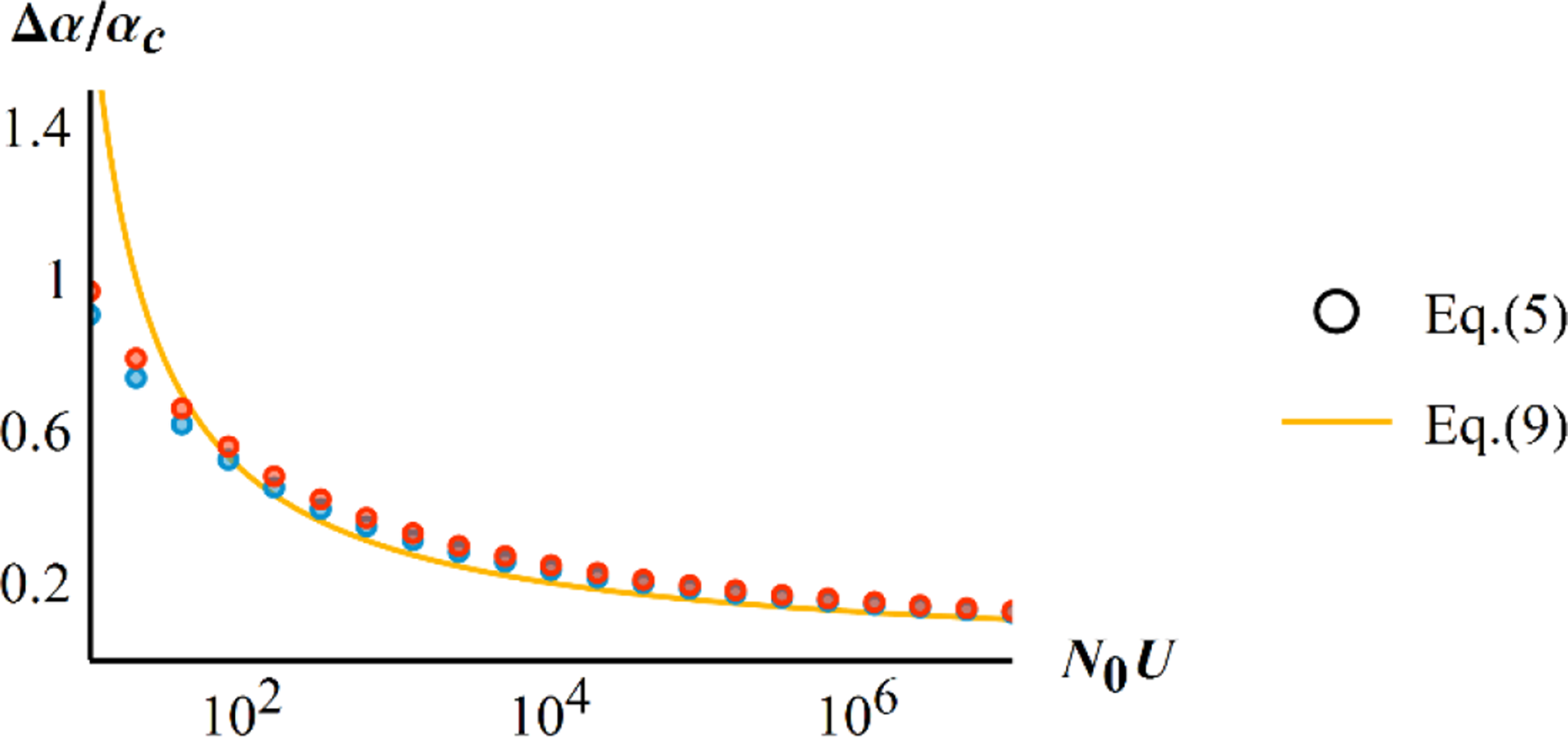
Scaled width of the characteristic stress window Δα/α_c_ versus the population-scale mutation rate *N*_0_*U*. The dots are obtained by numerical inversion of the ‘exact’ Eq.(5) with two values of r_max_ = 2 (blue) and r_max_ = 0.1 (red). The orange line shows the approximate scaled width of the characteristic stress window derived in Eq.(9). Other parameters are n = 4, λ = 0.005.

Interestingly, Eq.(9) provides a scale-free measure that may be compared across experiments, as it only depends on the genomic mutational input *N_0_U* (via *α*_c_). However, like all results so far, Eq.(9) only considers ER from *de novo* mutation. We now turn to ER from standing genetic variation.

### Rate of rescue from a population at mutation-selection balance

In the SSWM regime, and for a population at mutation-selection balance in the previous environment, each rescue event can be tracked back to either a pre-existing variant, or a *de novo* mutation (Orr and Unckless 2008; Martin *et al*. 2013; Orr and Unckless 2014). The proportion *ϕ_sv_* of rescue events caused by standing variance is then simply given by *ϕ_sv_ = ω_SV_/(ω_SV_ + ω_DN_*). A simple expression can be obtained again under the SME for *n ≥ 2* (see Eq.(A21) in the **Appendix**), but the approximation 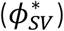 now further requires that decay rates are not vanishingly small 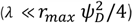.

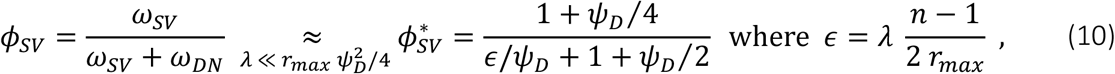

where ψ_*D*_ is defined in Eq.(6).

Eq.(10) captures the main features of how standing variance contributes to ER across (nonvanishing) stress levels (here, decay rate). Contrary to context-independent models, this contribution changes non-monotically with increasing stress level (**Fig.5**). At very mild decay rate *r_D_*, rescue relies on mild-effect mutations. The cost of such mutations - and hence their frequency before stress - is roughly independent of *r_D_* (Martin and Lenormand 2015), while their rate of production by *de novo* mutation decreases as *1/r_D_* (demographic effect), so the contribution of standing variance to ER increases with *r_D_* at small *r_D_*. In contrast at large stress levels, rescue stems from strong effect mutations. These mutations pay a substantial “incompressible cost” before stress that increases faster than *r_D_* (Martin and Lenormand 2015), while their rate of *de novo* production still decreases as *1/r*_D_, so the contribution of standing variance to ER decreases with *r_D_* at large *r_D_*. In the limit of very large *y_D_*, the distance between the two optima is very large and makes most of both the cost and decay rate, so that *c_H_(y*) ≈ *r_D_* for all mutations. Hence, *de novo* mutations and standing variants contribute equally to ER in this limit (*ϕ_S_v → 1/2)*.

**Figure 5:**
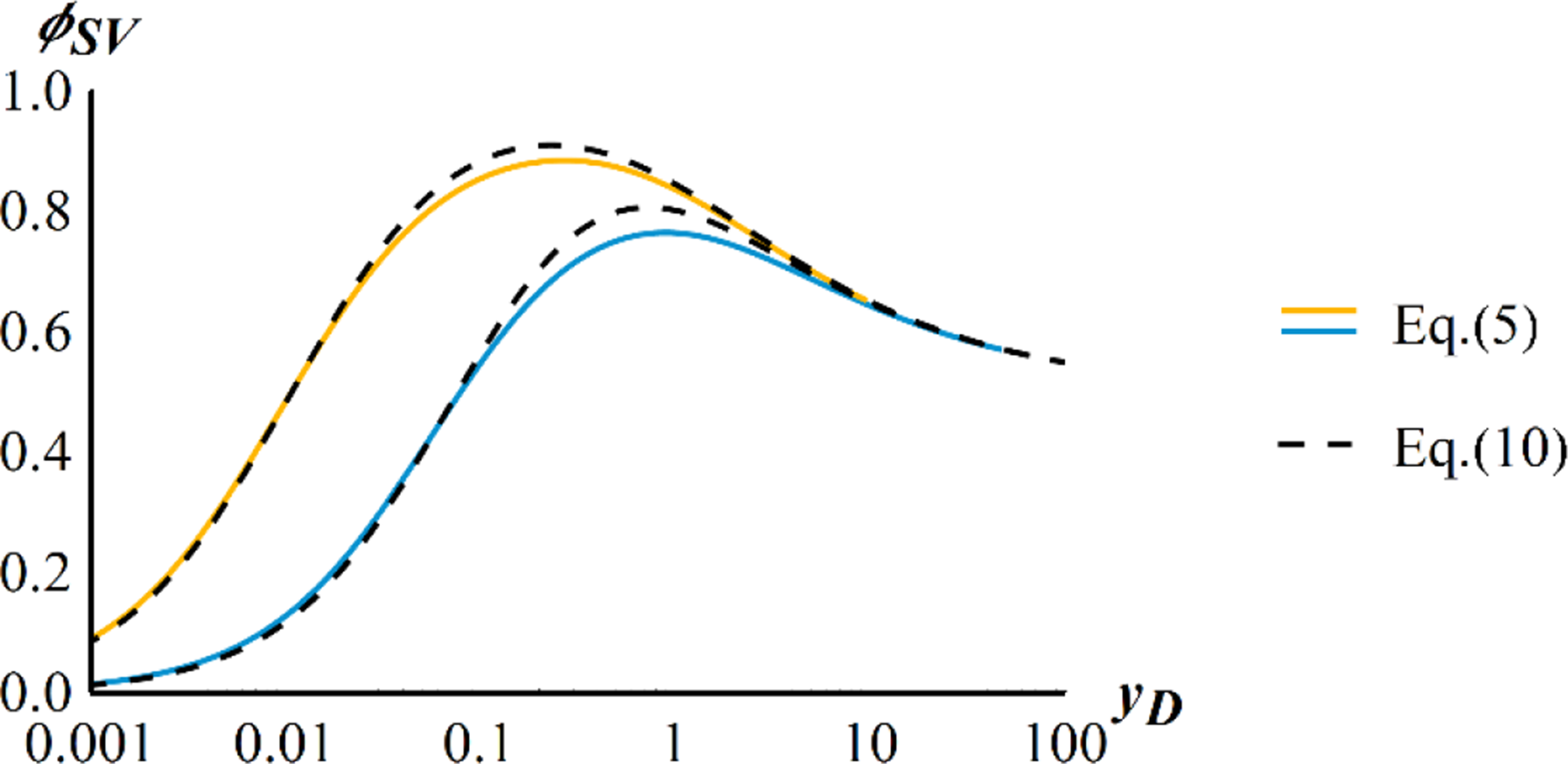
relative contribution of standing genetic variation to ER. The proportion of ER from standing variance is shown across scaled decay rates *y*_D_. The numerical computation for ϕ_SV_ (using Eq.(5) and (10)), for two values of r_max_ = 0.1 (blue) and r_max_ = 0.7 (orange), is compared to the approximate 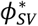 (Eq. (10), black dashed line). Other parameters as in **Fig.2.**

These different behaviors are illustrated in **Fig.5**, showing the variation of *ϕ_sv_* over a very wide range of scaled decay rates *y*_D_. The limit *ϕ_sv_* → 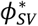 in Eq.(10) provides a fairly accurate approximation across the full range of stress levels. The limits when *y_D_ →* 0 and *y_D_ →* ∞ are in fact of limited biological interest, as they correspond to stress levels where ER becomes *de facto* certain or impossible, respectively. When focusing on the more biologically relevant range corresponding to the characteristic stress window, which occurs near the peak in *ϕ_sv_* in Fig.5 (see **Appendix Section IV subsection 4**), the variation of ***ϕ_sv_*** across stress levels becomes negligible. As illustrated in **Fig.6B**, *ϕ_sv_* remains close to 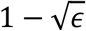 (see **Appendix** Eq.(A22)) as *y_D_* varies over a range where ER probabilities span several orders of magnitude (see **Appendix Section IV subsection 5**). Note that this behavior arises when stress only shifts the optimum (effect on *r_D_*), but does not affect peak height (*r_max_*) or the variance of mutational effects (*λ*).

**Figure 6:**
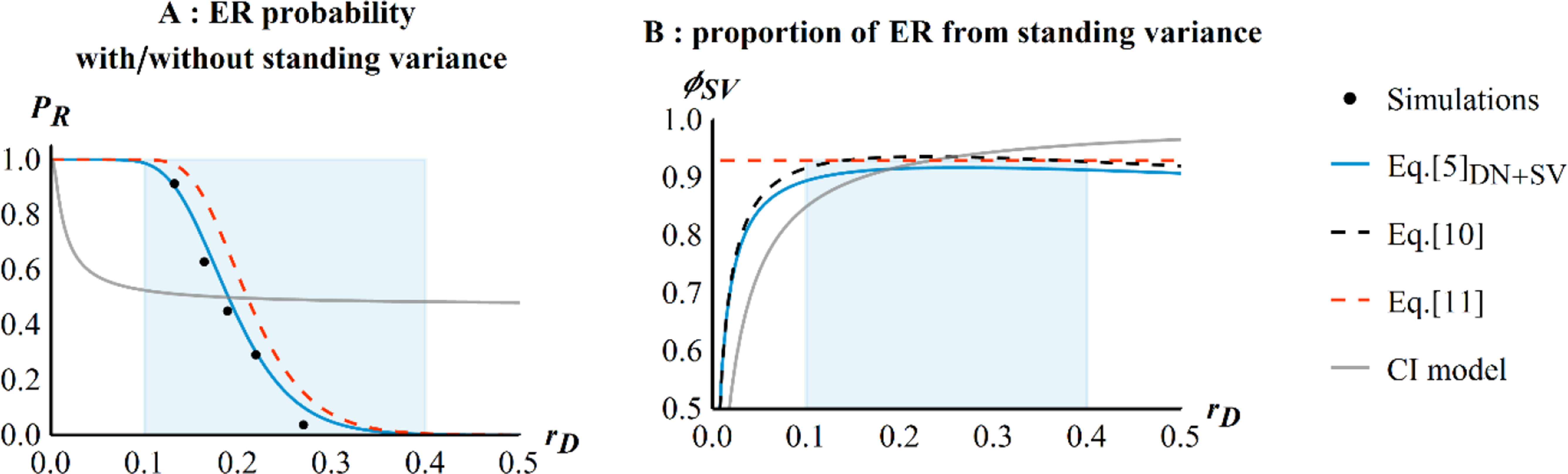
ER probability in the presence of standing genetic variation. In each panel, stress only affects the decay rate r_D_ (shifting optimum). In both panels, blue solid lines show the theory for *de novo* and standing variance (‘DN’+‘SV’) computed numerically (Eq.(5)) and the gray lines correspond to an equivalent theory without the FGM (named context-independent model as described in the Method section, “CI”) modified from in Orr and Unckless (2008). This last model was computed using a fixed proportion of resistant mutations equal to the one in Eq.(5) for a rescue probability of 0.5 (which explains why the two curves cross exactly at P_R_=0.5). The dashed red line gives the simpler expression for the overall rescue rate: 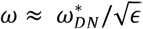 (Eq.(11)) with *∊* given in Eq.(10) and 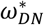 by Eq.(7). (A) ER probability in the presence of standing genetic variation as a function of r_D_ The dots give the results from simulations. (B) Proportion ϕ_SV_ of rescue from standing variance as a function of *r_D_* The black dashed-line give the approximate theory from Eq. (10) and the dashed red line max(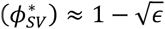) from Eq.(11). The shaded area shows the range of *r_D_* for which the ER probability drops from 0.99 to 10^−3^. Other parameters as in Fig.2.

Therefore the rate of rescue in the presence of standing variance is approximately proportional to that with only *de novo* mutation, with proportionality constant largely independent of the decay rate:

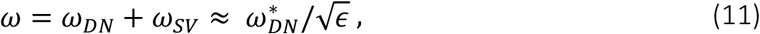

where *∊* is defined in Eq.(10). The rough constancy of *ϕ_sv_* also means that all the results obtained previously for ER from *de novo* mutations apply in the presence of standing variance, when stress only shifts the optima (as long as *n* ≥ 2).

The ER probability profile across stress levels (shown in **Fig.6A**) is the same as that from *de novo* mutations alone (**Fig.2**), but with a higher characteristic stress 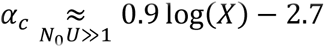from Eq. (8) with 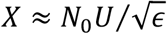). Moreover, when considering the contribution from standing variance, the difference between the FGM and context-independent models (gray line on **Fig.6A**) is striking. Indeed in the latter, the ER rate from standing variance (*ω_SV_*) is independent of the decay rate, hence *P_R_* saturates with stress to a constant value 1 – exp(− *N_0_ U q_R_/c_H_)*, where all ER events stem from standing variance.

Finally, note that when considering rescue from preexisting variance, *U* may change across environments, from *U_P_* (for previous) to *U_N_* (for new). For example, a stress-induced increase in DNA copy error would yield *U_N_ > U_P_*. Accounting for such shifts in mutation rate at the onset of stress, the total ER rate simply becomes 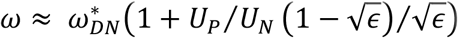 (from Eq.(11)).

## DISCUSSION

### Main results

We investigated the persistence of a population of asexual organisms under an abrupt environmental alteration. We assumed that this stress causes a shift in a multidimensional fitness landscape with a single peak (Fisher’s Geometrical Model - FGM -), which the population must ‘climb’ to avoid extinction. In such a landscape, faster population decline (due to stress-induced increase in the decay rate *r_D_*) necessarily means that resistance mutations are fewer, have lower growth rates in the presence of stress and higher costs in its absence. We believe that this constraint, not included in previous studies, adds a key element of realism to evolutionary rescue (ER) models. In our model, variation in stress levels may affect the landscape in various ways: shifting the optimum, changing the peak height, or altering the phenotypic scale of mutations (or the strength of stabilizing selection). Under a strong selection and weak mutation (SSWM) regime and assuming small mutational effects (SME), all these effects of stress on the distribution of mutation fitness effects are approximately captured by the variation, across stress levels, of a single composite parameter *α*, which is approximately independent of the dimensionality of the organism (number of orthogonal traits under selection). The probability of ER drops sharply with this effective stress level, more so than in previous ER models where the rate of population decline is decoupled from the input of resistant mutations. The characteristic stress window over which this drop occurs only depends on the initial population size *N_0_* and genomic rate of mutation *U*. As *N_0_U* gets large, the characteristic stress window reaches an asymptotic width (Δ*α* in Eq.(9)) while its center (the characteristic stress level *α_c_* in Eq.(8)) shifts towards higher values, approximately as log(*N_0_U)*.

When standing variance is available (population at mutation-selection balance before stress), its contribution to ER is dominant, and approximately constant across a wide range of stress levels that encompasses the characteristic stress window.

In Table 2, we summarize how these features compare to properties of previous ER models. We consider only the situation where stress shifts the position of the optimum, affecting *r*_D_ (as in previous models), because other effects of stress we investigate here (*r_max_* and *λ*) are not treated in previous models.

**Table 2:** Main results of “context-independent” models (Orr and Unckless 2008; Martin *et al.* 2013; Orr and Unckless 2014) and the present model (FGM) when stress only affects *r_D_* (only shifts the optimum position in our model). When the dependence to the parameters is given for the overall rate of rescue *ω*, it applies to both the rate of rescue from *de novo* mutations *ω_DN_* and from pre-existing variance *ω_SV_* (1) derived from the approximate expression 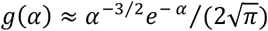.

|  | Context-independent model | FGM rescue model |
| --- | --- | --- |
| Impact of decay rate $r_D$ | $\omega_{DN} \propto r_D^{-1}$, $\omega_{SV} = \text{constant}$ | $\omega \propto e^{-r_D^2} r_D^{-3}$ (1) |
| Impact of mutational input $N_0 U$ | $\omega \propto N_0 U$ | $\omega \propto N_0 U$ |
| “Characteristic decay rate” ($r_D^c$) for DN rescue versus $N_0 U$ | $r_D^c \propto N_0 U$ | $r_D^c \propto \sqrt{\log(N_0 U)}$ |
| Relative contribution of standing variance to ER | Increases with $r_D$<br>$\phi_{SV} \approx \frac{r_D}{r_D + c}$ | ∼ Stable with $r_D$<br>$\phi_{SV} \approx 1 - \sqrt{\epsilon}$ |

### Genetic basis of ER patterns across environments

Our model allows identification of three ranges of stress levels that yield different eco-evolutionary patterns, despite all leading to extinction in the absence of evolution. First at low stress levels (*a* ≪ *a*_c_), although evolutionary change is required for persistence and demographic dynamics typical of ER may be observed (decay /rebounce), extinctions are *de facto* undetectable (ER is pervasive, *P_R_ ≈* 1). In this regime, we expect several resistance mutations to establish and co-segregate (frequent “soft sweeps” as in Wilson *et al*. 2017). Their number is predictable (≈ *N_0_ω*_R_), but the ultimate composition of the population in asexuals will depend on more complex clonal interference dynamics. Second, at intermediate stress levels (*α = 0(α_c_))*, small variation in stress conditions has large impact on the probability of population survival. Over this range, *P_R_ ≈ 1/2* so the expected overall number of rescue mutations in the population is less than one (*N_0_ω_R_ ≈* −log(*P_R_) = 0.7)*. Therefore, “hard sweeps” (including from standing variation) should be the most frequent: a single mutation typically establishes and rescues the population. Finally, at higher stress levels (*α ≫ a*_c_), very few populations overcome the imposed stress, and when they do it is typically through a hard sweep (*N_0_ω_R_* ≪ 1).

### Estimating parameters and testing the model

Studies on the emergence of resistance to controlled stress (e.g. antibiotics, fungicides, chemotherapy in cancer), especially in microbes, can generate a set of estimates of *P_R_* (the probability of resistance emergence), across stress levels. In general, to test (or use) predictions from ER models, it is critical to empirically relate physical measures of stress level (e.g. concentrations, temperatures, salinities, etc.) with demographic measures (e.g. decay rates). If we assume that the main effect of stress is to shift optimum positions, given a set of measurements of *r_D_* (“dose-kill curves”, Regoes *et al*. 2004), the change in ER probability with stress can be predicted via Eq.(7). This simple scenario of optimum shifting is the one that is considered by most of the literature on evolutionary ecology across environmental gradients, so it would seem natural to test it first. Furthermore, this scenario has received empirical support from an analysis of a few experimental studies of distribution of mutation effects on fitness across stress levels (Martin and Lenormand 2006b). However, these studies used mild stresses, which reduce growth without causing population decay. A more recent study on bacteria facing lethal doses of antibiotics, i.e. in the presence of decay (Harmand *et al*. 2017), suggests that factors other than the position of the optimum may also change with stress (*λ, r_max_, n)*. Estimating these extra parameters across environments can be challenging. The variance of mutational effects *λ* and the dimensionality *n* can be estimated by fitting the distributions of single random mutation effects on fitness (Martin and Lenormand 2006b; Perfeito *et al*. 2014), in a single environment if it is to be assumed constant, or in each environment otherwise. The maximal growth rate in the stress could be measured on lines well-adapted to the environment considered.

The effective stress level *α* is also amenable to empirical measurement and circumvents the issue of measuring joint changes in (*r_D_,r_max_,λ*) with stress. Consider a set of *P_R_* estimates across a range of empirically controlled stress levels, and some knowledge of the genomic (non-neutral) mutation rate (e.g. as estimated by mutation accumulation experiments) of the species and environment under study (*U)*. The initial population size *N_0_* is easily controlled by the experimenter. Then Eq.(7)*b*. suggests a simple estimator of *α* in each environment: *â = g^−1^(−log(1-P_R_)/N_0_U)*.

Finally, it is also possible to circumvent the problem of stress-induced variation in the parameters of the fitness landscape by considering multiple genetic backgrounds, in a single environment. Each background would have a given measurable decay rate *r*_D_, and other parameters (*λ,r_max_,n, U*) would be held fixed: while *λ, n* and *U* may change with the genetic background, this seems less likely than with the environment. The isotropic FGM assumes a strict equivalence between shifts in optima (multiple environments) from a given ancestor phenotype (single genetic background), and shifts in ancestor phenotypes (multiple genetic backgrounds) with a fixed optimum (single environment). The model could thus be applied and tested in this context. This could yield useful insights into the effect of epistasis (background dependence) on resistance emergence, an issue of notable importance when considering the fate of horizontally transferred resistance or multidrug resistance (as discussed in Wong 2017)

### Potential implications for resistance management

Our results suggest that stress levels have a strongly nonlinear impact on ER probabilities (**Fig.2**), at least in the context of abrupt environmental changes, in asexuals. This context may be particularly relevant to the chemical treatment of pathogens (cancer therapy, antivirals, antibiotics, fungicides, herbicides, etc.). In particular, the non-linear impact of stress on ER, if empirically confirmed, can provide insights regarding the optimization of treatment regimens or the quantification of the effect of poor treatment adherence on resistance emergence (e.g. in HIV, Harrigan *et al*. 2005). Our results point to the risk that even a slight lowering of drug doses (bellow prescribed treatment levels) could radically change the outcome of the treatment (making it *de facto* inefficient). On the contrary, a slight increase in prescribed doses could sometimes prove sufficient to allow efficient eradication.

## Limits and possible extensions

### Density-dependence and competitive release

Our model ignores density dependence, but some form may be easily introduced by considering a single density dependence coefficient (common to all genotypes) and using logistic diffusion approximations (Lambert 2005). This would potentially allow for “competitive release” (Read *et al*. 2011), whereby higher stresses may favor the emergence of resistance by rapidly depleting the sensitive wild-type population, thus releasing limiting resources for resistant genotypes. Previous models on competitive release assumed that the number of standing resistant mutants is independent of stress level (Read *et al*. 2011; Day and Read 2016). In this case, stress mostly limits *de novo* rescue mutation, with limited impact on the contribution from standing variance. On the contrary, the FGM imposes a similar drop, with stress, in the rate of rescue from *de novo* and preexisting mutants. The positive effects of competitive release on ER probability may thus be less important, in the FGM, than predicted from these previous models. Note however that more generally, the effect of density dependence on ER is more safely investigated by accounting explicitly for the effect of stress on the density-independent intrinsic rate of increase on the one hand (as done here in a density-independent model), versus on the competition component on growth (e.g. carrying capacity) on the other hand, than by compounding their effects into an overall density-dependent decay rate (e.g. Chevin and Lande 2010). This would require modeling the effect of stress on both the intrinsic rate of increase and the competition component, possibly through a landscape with two fitness functions, describing each component.

Anisotropy and parallel evolution in drug resistance: The present model is isotropic: all directions in phenotype space are equivalent (in terms of mutation and selection). In contrast, module-dependent anisotropy (where particular genes mutate along favored directions) can lead to substantial parallel evolution in the FGM (Chevin *et al*. 2010). Parallel evolution of resistance, whereby some (portions or sets of) genes contribute most of the resistance mutations, is often observed among drug-resistance alleles, and can increase with stress (Harmand *et al*. 2017), contrary to parallel evolution in a growing population, which is expected to decrease with increasing maladaptation (Chevin *et al*. 2010). Although not explored here, we conjecture that our model may accommodate mild anisotropy. Indeed, mild anisotropy (even environment-dependent) might have limited impact. If mutational covariances between traits merely “turn”, “shrink” or “expand” the phenotypic mutant cloud, this would approximately amount to a mere change in *λ* in an equivalent isotropic landscape (Martin and Lenormand 2006a; Martin 2014). However, a particular form of strong anisotropy may also arise where mutant phenotypes (in a given module) spread along a single favored direction (Martin 2014). Only this level of anisotropy would generate clear parallel evolution, and it will likely require implementing a fully anisotropic model.

### High mutation rates

Our results relied on a strong selection and weak-mutation (SSWM) approximation. When the mutation rate is higher (e.g. viruses or mutator bacterial strains), multiple mutants must be accounted for as a source of ER. These can in principle be introduced in the framework used here (Martin *et al*. 2013) but, especially when applied to the FGM, the results quickly become intractable. Alternative population genetics assumptions would then have to be used, but this is beyond the scope of the present work.

## Conclusion

Recently, the FGM has received renewed interest for its ability to provide testable, quantitative, and often accurate predictions regarding patterns of mutation effects on fitness, across various species and contexts (Tenaillon 2014). The present model is an attempt to extend its scope to model the evolution of resistance to stress. We hope that future experimental tests will evaluate its accuracy and potential to tackle various pressing applied issues.

## Acknowledgments

We thank the editor Joachim Hermisson and the reviewers for their help in improving this manuscript. In particular we thank Michael Kopp for extensive and very useful comments on our manuscript. This work benefited from discussions with S. Gandon, R. Gomulkiewicz, O. Tenaillon, F. Débarre and L. Roques. This work was supported by the French Ministry of Higher Education, Research and Innovation (MESRI allocation doctorale to Y.A.), Agence Nationale de la Recherche (ANR-13-ADAP-0016 “Silentadapt” to G.M. and ANR-13- ADAP-0006 “MeCC” to O.R.) and European Research Council (ERC-2015-STG-678140-FluctEvol to LMC). This is ISEM publication ISEM 2018-032.

